# Transcriptomic analyses reveal tissue-specific selection on genes related to apoptotic processes in the subterranean rodent, *Ctenomys sociabilis*

**DOI:** 10.1101/256875

**Authors:** Andrew Lang, Lauren Kordonowy, Eileen Lacey, Matthew MacManes

## Abstract

Specialization for a subterranean existence is expected to impact multiple aspects of an organism’s biology, including behavior, physiology, and genomic structure. While the phenotypic correlates of life underground have been extensively characterized, the genetic bases for these traits are not well understood, due in part to the challenges of generating large, multi-locus data sets using traditional DNA sequencing strategies. To begin exploring the genomic architecture of adaptation to a subterranean existence, we generated high-quality *de novo* transcriptome assemblies for 8 different tissue types (hippocampus, hypothalamus, kidney, liver, spleen, ovary, testis, skin) obtained from the colonial tuco-tuco (*Ctenomys sociabilis*), a group-living species of subterranean rodent that is endemic to southwestern Argentina. From these transcriptomes, we identified genes that are evolving more rapidly in the *C. sociabilis* lineage compared to other subterranean species of rodents. These comparisons suggest that genes associated with immune response, cell-cycle regulation, and heavy metal detoxification have been subject to positive selection in *C. sociabilis*. Comparisons of transcripts from different tissues suggest that the spleen and liver - organs involved in immune function and detoxification - may be particularly important sites for these adaptations, thereby underscoring the importance of including multiple tissue types in analyses of transcriptomic variation. In addition to providing an important resource for future genomic studies of *C. sociabilis*, our analyses generate new insights into the genomic architecture of functionally significant phenotypic traits in free-living mammals.

## INTRODUCTION

Convergent traits provide critical opportunities to examine interactions between shared environmental challenges, selection, and the evolution of phenotypic and genotypic variation (Mares, 1975; Muschick, Indermaur & Salzburger, 2012; Parker et al., 2013). One well-characterized suite of convergent phenotypic traits occurs among subterranean rodents, which are defined by their tendency to spend virtually their entire lives in underground burrows (Nevo, 1979; Lacey & Patton, 2000). This designation encompasses more than 120 species representing 6 families and 3 suborders of rodents (Lacey & Patton, 2000; Gardner, Wilson & Reeder, 2005). Shared physiological challenges associated with life underground include the high energetic costs of excavating burrows (Luna & Antinuchi, 2006; Zelová et al., 2011), hypoxia and hypercapnia (Lovegrove, 1986; Buffenstein, 2000), maintenance of circadian patterns of activity (Vasicek et al., 2005; Urrejola et al., 2005; Tomotani et al., 2012), and, at least in some habitats, exposure to heavy metals in soils (Lovegrove, 1986; De Vleeschouwer et al., 2014; Fernández-Cadena et al., 2014). While the convergent phenotypic processes associated with these challenges have been studied in some detail (Nevo, 1979; Buffenstein, 2000; Burda, Šumbera & Begall, 2007), the genetic architecture underlying similar physiological responses to these challenges remains largely unknown (but see Partha et al., 2017). Determining the proximate mechanisms (*e.g*., the genetic underpinnings) of adaptations enabling organisms to thrive in such an environment is critical to improving our understanding of how specializations for a subterranean existence arise and are maintained.

The advent of high-throughput transcriptome sequencing has greatly facilitated efforts to relate patterns of gene expression to differences in phenotypic traits, including physiological processes such as metabolism (Devi et al., 2016) and water regulation (Kordonowy & MacManes, 2016; MacManes, 2017). This sequencing strategy has also been used to identify physiologically relevant regions of the genome undergoing positive selection (Zhang, Dyer & Rosenberg, 2000; Swanson et al., 2001; Brodsky et al., 2005; Kosiol et al., 2008; Karn et al., 2008; Gardiner et al., 2008; Kong et al., 2011), thereby generating insights into the evolutionary bases for relationships between gene expression and specialization for specific phenotypic attributes. The marked examples of evolutionary convergence and divergence among burrow-dwelling mammal species offer an ideal opportunity to implement sequencing methods for exploration of the genomic bases for the functional and evolutionary consequences of a shared subterranean lifestyle.

The colonial tuco-tuco *(Ctenomys sociabilis*) is a subterranean rodent that is endemic to Neuquen Province, Argentina (Tammone, Lacey & Relva, 2012). This species has been the subject of extensive research due to its unusual social system; while the majority of ctenomyids are thought to be solitary, *C. sociabilis* is group living, with burrow systems routinely occupied by multiple adult females plus, in many cases, a single adult male (Lacey, Braude & Wieczorek, 1997; Lacey & Wieczorek, 2004). In particular, this species has been studied with respect to not only to behavior, ecology and demography (Lacey, Braude & Wieczorek, 1997; Lacey, 2001; Chan & Hadly, 2011; MacManes & Lacey, 2012), but also neuroendocrinology (Beery, Lacey & Francis, 2008; Woodruff et al., 2013) and population genetic structure (Lacey, 2001; Hambuch & Lacey, 2002; Chan et al., 2005). Compared to other subterranean rodents for which transcriptomic data are available (Malik et al., 2011; Lin et al., 2014), *C. sociabilis* is phylogenetically, geographically, and behaviorally distinct, suggesting that this species is critical to efforts to examine the genomic impacts of adaptation to life underground.

Here we present a high-quality annotated transcriptome generated from eight tissue types (hippocampus, hypothalamus, kidney, liver, spleen, ovary, testis, skin) obtained from *C. sociabilis*. The use of multiple tissues has resulted in a particularly complete transcriptome for htis non-traditional study species. In addition to presenting this annotated assembly, we characterize each tissue type with regard to the most highly abundant transcripts, after which we compare patterns of expression across tissue types. We then conduct a comparative analysis of coding sequence evolution in *C. sociabilis* based on contrasts with single-tissue transcriptomes from seven other subterranean rodent species. In addition to highlighting the importance of tissue type in determining patterns of transcript abundance, our analyses generate important new insights into the genetic correlates of subterranean life.

## METHODS

### Sample collection, RNA Extraction & Library Preparation

Tissue samples were obtained from two adult *C. sociabilis* (1 male and 1 female) that were members of a captive population of this species maintained at the University of California, Berkeley. The housing and husbandry of this population have been described previously (MacManes & Lacey, 2012; Woodruff et al., 2013). The animals sampled were euthanized via overdose with Isoflurane followed by decapitation. The hippocampus, hypothalamus, kidney, liver, ovary, skin, and spleen were extracted from the female and the testes were extracted from the male. Each tissue type was placed in a cryotube containing RNAlater (Thermo Fisher Scientific, Waltham, MA) and then flash frozen with liquid nitrogen. The interval between euthanasia and flash freezing of tissues did not exceed five minutes. All tissue samples were stored at −80°C until they were sent to the Broad Institute (Cambridge, MA) for RNA extraction, cDNA library preparation, and 125bp paired-end sequencing on an Illumina 2500 platform. All procedures involving live animals were approved by the Berkeley Animal Care and Use Committee and were consistent with guidelines established by the American Society of Mammalogy for the use of wild mammals in research (Sikes, 2016).

### Tissue-Specific Transcriptome Assembly

Tissue-specific Illumina reads (36-49 million paired-end reads per tissue) were obtained for each of the 8 tissue types examined. For each tissue type, read quality was evaluated with SolexaQA++ v3.1.4 (Cox, Peterson & Biggs, 2010) and reads were corrected using Rcorrector v1.0.1 (Song & Florea, 2015). Adaptor sequences and reads falling below the quality threshold PHRED=2 were removed using Trimmomatic (Bolger, Lohse & Usadel, 2014), following the protocol of MacManes (2014). *De novo* transcriptome assemblies were generated using Trinity v2.1.1 (Haas et al., 2013). For each tissue type, two assemblies were generated – a khmer normalized (Crusoe et al., 2015) 100x coverage assembly and a non-normalized assembly. Digital normalization had no detectable effect on either the completeness or the consistency of the resulting transcriptomes (Table S1) and thus all downstream analyses were conducted using assemblies generated from the non-normalized datasets.

### Compiled Transcriptome Assembly, Annotation and Analysis

In addition to tissue-specific transcriptomes, read data from all tissue types were pooled to generate a single, merged transcriptome assembly. We produced 12 alternative merged assemblies through combinations of read subsampling, transrate optimization, and merging algorithms. Each assembly was evaluated for quality using TransRate v1.0.1 (Smith-Unna et al., 2016), which generates both quality metrics and an optimized assembly. In addition, we evaluated each assembly for completeness using the Vertebrata database within BUSCO v1.1b1 (Simão et al., 2015). Based on these analyses, we selected the assembly with the highest quality and completeness. The pipeline for producing this selected assembly is described below. Because previous research has revealed that little information is gained from using datasets above 40M reads (MacManes, 2015), a random subset of 50 million paired-end reads were selected for analyses (seqtk v1.0-r82 (https://github.com/lh3/seqtk)) from the entire dataset (N= 339 million reads). The subset of reads was assembled with both Trinity v2.1.1 and BinPacker (Liu et al., 2016). The resulting two assemblies were merged into a single assembly using Transfuse v0.5.0 (https://github.com/cboursnell/transfuse). This merged assembly was optimized with TransRate to retain only highly-supported contigs. The resulting assembly was annotated with dammit version 0.3.2 (https://github.com/camillescott/dammit) and filtered to retain only annotated transcripts.

### Transcript Abundance and Gene Presence/Absence

To generate measures of relative transcript composition across tissue types, the abundance of each annotated transcript in our tissue-specific assemblies was assessed using Kallisto v0.42.4 (Bray et al., 2016). Transcripts with TPM (transcripts-per-million) values of less than 1 were determined to be absent from a given tissue (MacManes et al., 2017; MacManes & Lacey, 2012; MacManes & Eisen, 2014). Transcript presence/absence was compared across all tissues using the UpSetR package (Gehlenborg, 2016), and the 10 most abundant genes were identified within each tissue.

### Comparative Analysis with Other Subterranean Taxa

To compare patterns of gene evolution across multiple lineages of subterranean rodents, we downloaded Illumina RNAseq reads for 5 other subterranean species (*Spalax carmeli*, *Bathyergus suillus*, *Tachyoryctes splendens*, *Eospalax baileyi*, *Cryptomys hottentotus pretorian*) from the NCBI Sequence Read Archive (accession numbers SRR2016467, SRR2141210, SRR214121, SRR931783, and SRR2141213, respectively). In addition, we downloaded mRNA datasets derived from whole genome sequencing projects for a sixth species of subterranean rodent (*Heterocephalus glaber:* Mole Rat genome v1.7.2 http://gigadb.org/dataset/100022) and for *Mus musculus (Mus* genome vGRCm38); the latter served as the outgroup for these analyses. These mRNA data sets were assembled following the Oyster River Protocol (http://oyster-river-protocol.readthedocs.io/, (MacManes, 2015)). Together with the transcripts for *C. sociabilis* generated here, this comparative data set encompassed 3 families of subterranean rodents (Ctenomyidae, Spalacidae, Bathyergidae), each of which represents a phylogenetically distinct origin of specialization for life in underground burrows.

For each of the species in this comparative data set, coding sequences were identified using TransDecoder v3.0.0 (Haas et al., 2013). Orthologous relationships among these species (including the *M. musculus* outgroup) were identified using the output from BUSCO v2.0 and the associated database of mammalian sequences. The resulting groups of orthologous transcripts were then edited to include only single copy transcripts, which were then aligned using Prank v150803. Sequence alignments were refined using pal2nal v14 (Suyama, Torrents & Bork, 2006) and a gene tree was constructed using RAxML v8.2.8 (Stamatakis, 2014). To explore potential evidence of selection on the genes included in our dataset, we used PAML v4.9a (Yang, 2007), with our gene tree as the phylogenetic framework. Specifically, we tested for positive selection using the M7 versus M8 models in PAML. We then tested for evidence of lineage-specific selection using the PAML branch-site model with *C. sociabilis* as the foreground lineage. We controlled the false discovery rate for multiple comparisons following the procedure of Benjamini and Hockberg (1995). Genes determined to be under positive selection were then examined using the Gene Ontology Consortium Enrichment Analysis (http://geneontology.org/page/go-enrichment-analysis) tool to determine if these loci were grouped according to ontology terms.

To explore potential tissue-specific patterns of gene expression among loci identified as being under positive selection in *C. sociabilis*, we imported gene expression count data generated by Kallisto into the R statistical package v3.3.0 (Team, R C, 2013). To allow comparisons across tissue types, we normalized count data using the TMM method (McCarthy, Chen & Smyth, 2012) as implemented in edgeR v3.1.4 (Robinson, McCarthy & Smyth, 2010). For each transcript under positive selection, we identified the tissue for which the expression level was highest. These maximum count values were then normalized by dividing by the total number of genes expressed in that tissue; this procedure allowed us to identify tissues enriched for positively selected transcripts.

### Data and Code Availability

Sequence read files for this study are available on the NCBI Short Read Archive (PRJNA358281). All code used in transcriptome assembly, annotation, analyses, and data visualization is freely available online at (https://github.com/macmanes-lab/tuco_manuscript and https://github.com/macmanes-lab/paml). The tissue-specific assemblies, as well as the final merged *C. sociabilis* transcriptome assembly are available on Dropbox (in fasta format), as are all annotation data files (in gff3 format) and kallisto transcript counts (https://www.dropbox.com/sh/jq98iderelxi9sm/AAAQG6Ex51sG9dcIrb8vK8gPa?dl=0). These files will be uploaded to Dryad upon acceptance of this manuscript for publication.

## RESULTS AND DISCUSSION

### Tissue-specific Transcriptome Assembly Analysis

Individual tissue-specific transcriptome assemblies were 68-82% complete (mean= 75.87%), with TransRate scores ranging from 0.145 to 0.172 (Table S1). The TransRate optimized assemblies, which included only highly-supported transcripts, contained on average 7% fewer BUSCOs than the original assemblies. Due to this pronounced reduction in completeness, the TransRate optimized assemblies were not used for subsequent analyses. While individual, non-optimized tissue-specific assemblies were of acceptable quality and completeness, they were notably inferior in quality and completeness to the compiled, transfused assembly described below.

### Compiled Transcriptome Assembly, Annotation and Analysis

The most complete and highest quality assembly was generated from a 50 million read-pair subsample of the full dataset (Table 1). This assembly was annotated and all non-annotated transcripts were removed to produce the final assembly (annotation_only; Table 1). Removal of unannotated transcripts resulted in minimal reduction of TransRate and BUSCO scores but reduced the number of contigs by ~ 50%; the transcripts removed were likely artifacts of the assembly process (Moreton, Izquierdo & Emes, 2016) and thus this reduction was was not considered problematic. Reads from different tissue types mapped to the final transcriptome at a rate of 86-90% (Table 2). The final assembly contained 96,224 annotated transcripts, with 79,938 search matches to the Uniref90 database, 73,896 matches to OrthoDB (Waterhouse et al., 2013), 46,659 matches to PFAM, and 2,698 matches to RFAM (Griffiths-Jones et al., 2005). Of the 96,224 transcripts in this final assembly, 78,241 (81.3%) contained open reading frames (ORFs) and 53,711 (55.8%) contained complete ORFs, indicating that these transcripts included the entire protein-coding sequence for the associated locus.

**Table 1.** A comparison of assemblies utilizing metrics for quality and completeness. (Num. Reads = Number of Reads, Num. Contigs = Number of Contigs, Assembly size, TransRate score, and BUSCO Metrics: C = Complete, D = Duplicated, M = Missing BUSCOs). The good_compiled_50M_transfuse assembly was chosen for annotation, and the annotation_only assembly is the transcriptome we present as our finalized assembly.

| Assembly | Num. Reads | Num. Contigs | Assembly Size | Transrate Score | BUSCO Metrics |
| --- | --- | --- | --- | --- | --- |
| good_compiled_50M_transfuse | 50M | 157996 | 240Mb | 0.430 | C: 88%, D: 64%, M: 8% |
| annotation_only | 50M | 96224 | 227Mb | 0.420 | C: 88%, D: 64%, M: 8% |

**Table 2.** Burrows-Wheeler Aligner mapping statistics comparing the percent mapping and percent properly paired mapping rates of the annotated assembly (annotated good_compiled_50M_transfuse) and the final assembly (Annotation.only).

| Tissue | good_compiled_50M_transfuse |  | Annotation_only |  |
| --- | --- | --- | --- | --- |
|  | % mapped | % prop paired | % mapped | % prop paired |
| hippocampus | 87.51 | 80.28 | 85.18 | 78.51 |
| hypothalamus | 86.40 | 79.10 | 83.98 | 77.29 |
| kidney | 86.74 | 77.05 | 84.53 | 75.47 |
| liver | 87.79 | 79.68 | 85.92 | 78.50 |
| ovary | 84.59 | 74.62 | 81.57 | 72.42 |
| testes | 85.24 | 77.00 | 82.86 | 75.32 |
| skin | 85.82 | 73.42 | 83.37 | 71.71 |
| spleen | 90.67 | 83.53 | 88.89 | 82.24 |
| Average | 86.85 | 78.09 | 84.54 | 76.43 |

### Comparative analysis with Other Subterranean Taxa

Using the output from BUSCO, we identified 2,182 single-copy ortholog groups from the transcriptomes of seven subterranean rodent species and from *Mus musculus*. Of these, 1,951 (89.4%) were successfully aligned and analyzed via PAML software. Branch site analysis identified 50 transcripts as being under positive selection in the lineage leading to *C. sociabilis;* in contrast, only seven were identified using the site-model of positive selection. While the larger set of transcripts identified using the branch-site model for GO enrichment did not reveal statistically significant enrichment of GO terms for *C. sociabilis* genes under positive selection, it did reveal that many of the GO terms identified corresponded to processes of cell proliferation control, DNA damage response, immune response, and ion transport. These findings are intriguing in light of evidence suggesting that burrowing rodents may be exposed to heavy metals or other toxins in the soils that they inhabit (De Vleeschouwer et al., 2014; Fernández-Cadena et al., 2014) and recent studies characterizing the immunogenetics of subterranean rodents (Cutrera et al., 2010; Merlo, Cutrera & Zenuto, 2016; Novikov et al., 2016). Particularly exciting is the identification of transcripts involved in the control of cell proliferation, which has potential ties to susceptibility to cancer (Tian et al., 2013).

For each gene under positive selection, we identified the tissue in which it was most abundant (Figure 1). We then compared the number of positively selected genes per tissue to that expected under a random distribution of these loci across tissue types – that is we divided the 50 genes under positive selection by the number of tissues (N=8) sequenced and then normalized these values according to the overall number of genes expressed in each tissue. This analysis revealed a significantly higher representation of genes under positive selection in the spleen and liver (*χ*^*2*^ test, p-values <0.05), an outcome that is perhaps not surprising given the functional roles of these tissues. Collectively, the preponderance of genes under positive selection in *C. sociabilis* that are associated with response to cell damage and immune response suggests that the environmental physiology of this species deserves further investigation.

**Figure 1.**
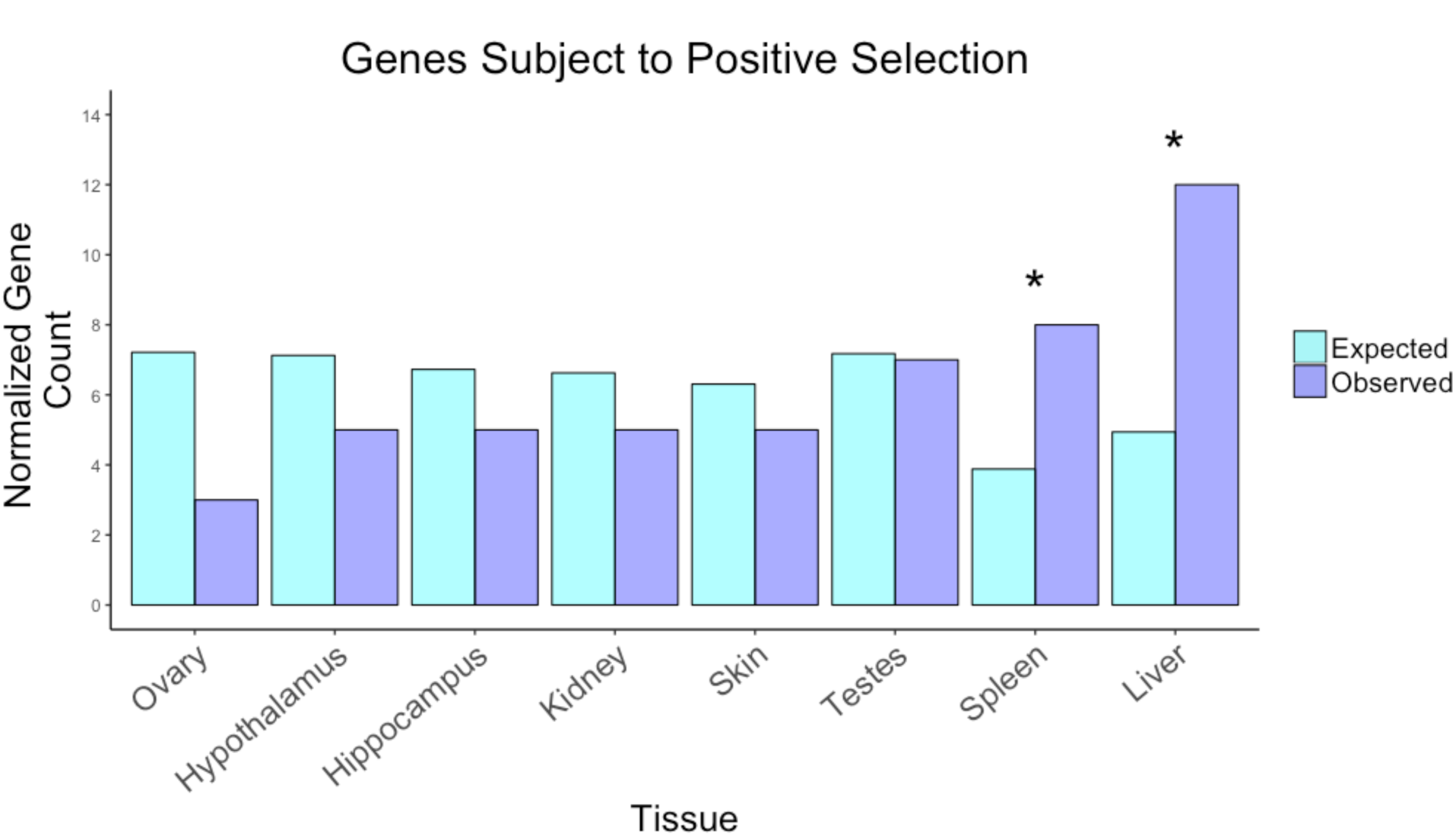
Tissue-specific counts of the 50 positively selected genes detected, normalized by the total number of genes present in each tissue. Tissue types are indicated on the x-axis. Expected abundance of positively selected genes is depicted by light blue bars; observed abundance of positively selected genes is shown in dark blue. Asterisks denote statistically significant differences between expected and observed values (Chi-square tests, p < 0.05).

### Transcript Abundance and Gene Presence/Absence

Filtering of transcripts to remove those for which TPM was less than 1 (Havens & MacManes, 2016; Kordonowy & MacManes, 2016) removed 5,722 (6.0%) of our annotated transcripts. Of the remaining 90,502 transcripts, 21,602 (23.9%) were expressed in all of the tissue types examined. In contrast, 774 (0.9%) of these transcripts were expressed in only a single tissue type. The distribution of these unique transcripts across tissue types was as follows: skin (N = 171), liver (N = 156), testes (N = 140), ovary (N = 93), spleen (N = 92), kidney (N = 77), hypothalamus (N = 23), and hippocampus (N = 22).

Between 81% and 88% of reads mapped to the reference transcriptome. Visual representations of transcript overlap between tissue types are presented in Figures 2, S1, and S2. The 10 most common transcripts unique to each tissue type are shown in Figure 3. While our data set did not allow a statistical comparison of levels of gene expression across tissue types, our assessments of transcript abundance per tissue type provide potential insights into the function of each tissue examined (Table S2). In particular, pairwise comparisons of transcript abundance revealed that tissues with similar functions tended to display similar suites of highly-expressed transcripts. For example, the two brain tissues examined – the hippocampus and the hypothalamus – shared the highest number of transcripts (5,200 out of 62,716 and 66,421 transcripts, respectively). The two reproductive tissues examined – the testes and the ovary – had an overlap of 1,359 out of 66,876 and 67,251 transcripts, respectively. The spleen did not share many transcripts with other tissues; the greatest overlap in spleen transcripts was with the testes (400 of 66,876 transcripts) and the ovary (298 of 67,251 transcripts). The kidney and liver, both associated with detoxification, shared 1,382 of 61,767 and 46,063 transcripts, respectively. Somewhat surprisingly, of the 58,796 transcripts in the skin, this tissue shared 1,397 with the ovary, the largest transcript overlap of any other tissue with the skin.

**Figure 2.**
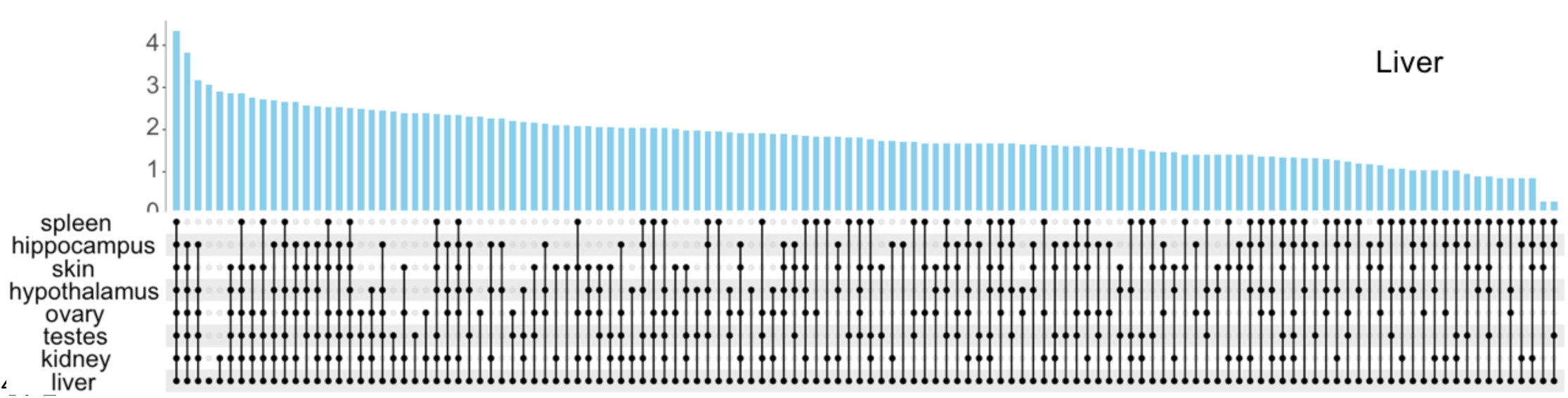
Comparing transcript composition of the liver to other tissues. The x-axis depicts intersections between tissue types, and the y-axis is the log_10_ transformation of normalized transcript counts. The 128 Intersection groups have been arranged to present groups with the highest transcript counts to the left, and lowest counts to the right. Figures depicting transcript composition of the remaining tissues can be found in supplemental materials (Figures S1 & S2).

**Figure 3.**
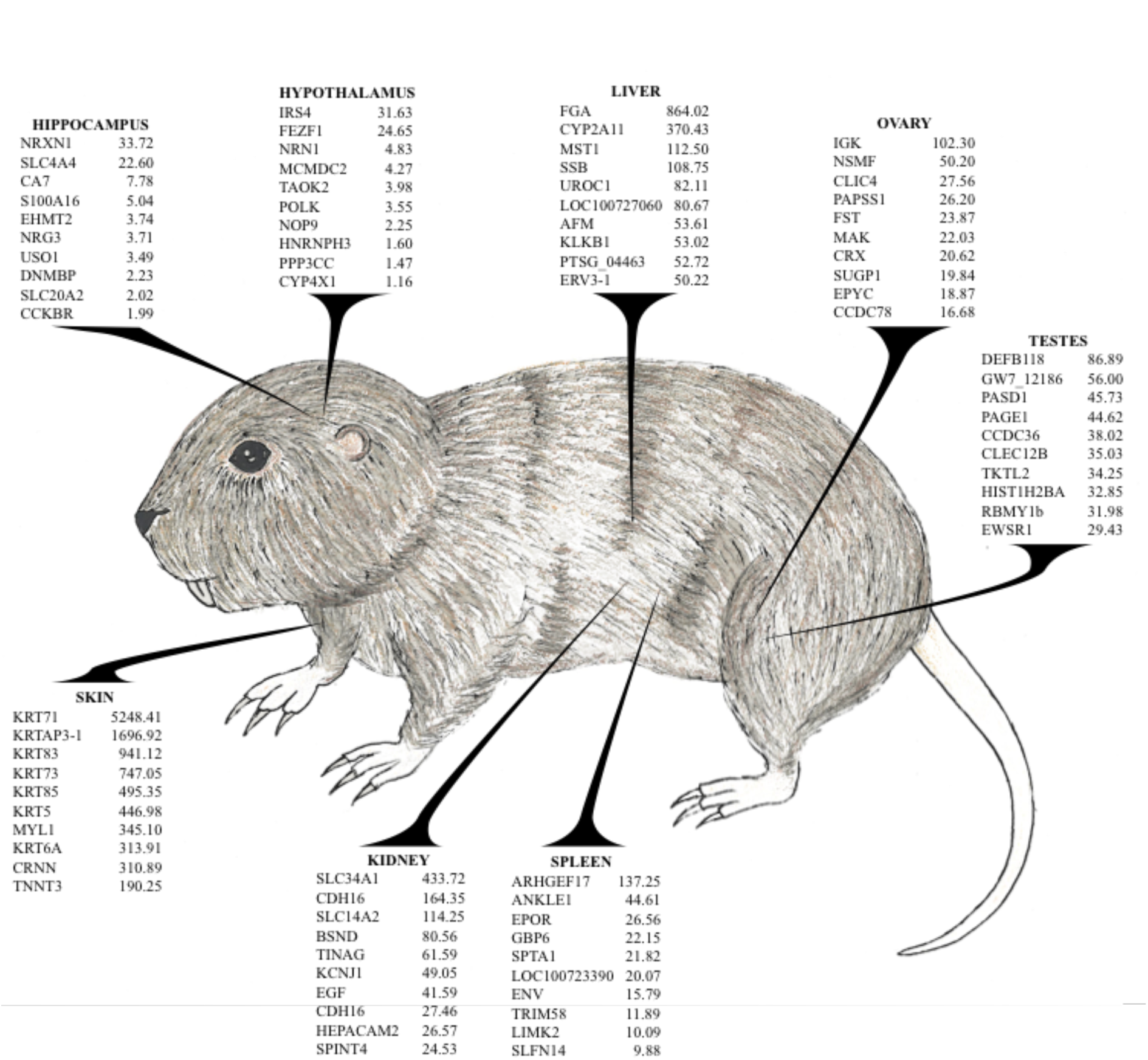
Ten most abundant unique transcripts for each tissue type. For each tissue type, the left column is the gene ID, while the right column contains the associated TPM values.

### Tissue Characterization

Each of the tissue types included in this study has been well characterized with respect to its function in mammalian biology. Accordingly, we examined whether functional differences between tissues were reflected in the identities of the most abundant transcripts unique to each tissue. We also assessed loci under positive selection, highlighting aspects we believe may be key factors associated with live in underground burrows. The functions of many of the most abundant transcripts that were unique to a given tissue type have been characterized as part of empirical studies, as described below:

#### The hippocampus

The hippocampus is integrally involved in neurotransmission (Vianna et al., 2000; Shatz, 2009). In particular, the hippocampus has been studied with regard to spatial memory and navigation (Bannerman et al., 2002; Eichenbaum, 2017) and as a site for for adult neurogenesis in the mammalian brain (Seri et al., 2001; van Praag et al., 2002). Among the transcripts that were uniquely abundant in the hippocampus in *C. sociabilis* were genes associated with regulating presynaptic density (Neurexin: NRXN1, TPM= 33.72) and synchronous firing of hippocampal pyramidal cells (Carbonic Anhydrase VII: CA7, TPM= 7.78) (Ruusuvuori et al., 2004; Kumar & Thakur, 2015). Loci found to be under positive selection in the hippocampus include genes involved in cell cycle progression and tumor growth, such as BRCA1 Associated Protein 1 (BAP1) and Apoptosis Antagonizing Transcription Factor (AATF) (Bruno et al., 2002; Qin et al., 2015). Both of these genes have been implicated in tumor suppression and cell growth inhibition, with BAP1 functioning by means of deubiquitinating host cell factor-1 (Machida et al., 2009) and AATF acting as an essential cofactor for the p53 gene (Bruno, Iezzi & Fanciulli, 2016).

#### The hypothalamus

The hypothalamus has been implicated in multiple critical signaling pathways, such as the Hypothalamic-Pituitary-Adrenal (stress) and Hypothalamic-Pituitary-Gonadal (reproductive) axes in vertebrates (Hall et al., 2012; Clément, 2016). Transcripts that were uniquely abundant in the hypothalamus tended to be directly involved in downstream signaling of activities such as feeding and parental or sexual behaviors (Insulin Receptor Substrate 4: IRS4, TPM= 31.63) as well as formation of the diencephalon and prethalamic brain region (FEZ Family Zinc Finger 1: FEZF1, TPM= 24.65) (Numan & Russell, 1999; Shimizu & Hibi, 2009). Genes identified to be under positive selection, similar to those identified for the hippocampus, are implicated in the cell cycle. For example, Prostate Androgen-Regulated Mucin-Like Protein 1 (PARM1) functions in prostate cell androgen dependence, has been linked to apoptotic mechanisms (Bruyninx et al., 1999), and may impart cell immortalisation (Cornet et al., 2003).

#### The ovary

Ovarian function is highly regulated by hormonal signals that mediate cell proliferation and the production of viable ova (Verga Falzacappa et al., 2009). Transcripts that were uniquely abundant in the ovary included an immunogene (Immunoglobulin Kappa Locus: IGK, TPM= 102.30) as well as genes involved in neuron development (NSMF, TPM= 50.20), and primordial follicle formation (Follistatin: FST, TPM= 23.87) (Brekke & Garrard, 2004; Palevitch et al., 2009; Kimura, Bonomi & Schneyer, 2011). Ovarian genes under positive selection (e.g., Nuclear Mitotic Apparatus Protein 1; NUMA1) tend to function in the structural components of cellular division and mRNA binding. For example, Nuclear Mitotic Apparatus Protein 1 (NUMA1) interacts with proto-oncogene PIM1 during mitosis and regulates p53-mediated transcription (Bhattacharya et al., 2002).

#### The testis

Similar to the ovaries, testis function is regulated hormonally and results in the production of viable gametes (Alves et al., 2013; O’Shaughnessy, 2014). The uniquely most abundant testis transcripts included an antimicrobial defense immunogene (Beta-defensin: DEFB118, TPM= 86.89), a transcription factor (PAS Domain Containing 1: PASD1, TPM= 45.73), and a gene unique to the testes that has not been fully characterized with regard to structure or function (P Antigen Family, Member 1: PAGE1, TPM= 44.62). Testicular genes under positive selection include known regulators of DNA damage (SprT-Like N-Terminal Domain; SPRTN, Ring Finger and WD Repeat Domain 3; RFWD3) (Fu et al., 2010; Gong & Chen, 2011; Liu et al., 2011; Juhasz et al., 2012) and cell proliferation regulation (Dishevelled Segment Polarity Protein 3, DVL3) (Schlange et al., 2007). Thus, as in the ovary, active testicular genes were generally associated with immune response and cell replication.

#### The skin

Not surprisingly, the majority of the most abundant transcripts that were uniquely abundant in skin were keratins (Keratin 71 Type II: KRT71 TPM= 5248.41, Keratin Associated Protein 3-1 Type II: KRTAP3-1 TPM= 1696.92, Kerain 83: KRT83 TPM= 941.12, Keratin 73 Type II: KRT73 TPM= 747.05, Keratin 85 Type II: KRT85 TPM= 495.35, Keratin type II cytoskeletal 5: KRT5 TPM= 446.98), the proteins that comprise the protective external layer for epithelial cells (Bragulla & Homberger, 2009; Deek et al., 2016). Highly abundant skin transcripts also include genes involved in muscle movement (Myosin Light Chain 1; MYL1 TPM= 345.10, Troponin T3; TNNT3 TPM= 190.25) (Periasamy et al., 1984; Ling et al., 2010; Wei & Jin, 2016). Genes found to be under positive selection in skin have been associated with tumor suppression (UBS Domain Protein 1; UBXN1) (Wu-Baer, Ludwig & Baer, 2010) and repair of double-stranded DNA (Heterogeneous Nuclear Ribonucleoprotein U Like 1 (HNRNPUL1) (Polo et al., 2012).

#### The kidney

Two well-documented functions of renal tissue are the transport of nutrients and the secretion of urine (Wang & Giebisch, 2009; Bobulescu & Moe, 2012). Consistent with this, uniquely abundant transcripts identified in the kidney included solute carriers SLC34A1 (TPM= 433.72) and SLC14A2 (TPM= 114.25), which are involved in transport of nutrients and urea (Shayakul, Clémençon & Hediger, 2013; Martovetsky, Bush & Nigam, 2016). Among those genes displaying signatures of positive selection in the kidney were Suppressor of Ty 3 (SUPT3), which binds p53 during DNA repair (Martinez et al., 2001; Gamper & Roeder, 2008) and N-Myc Downstream Regulated 1 (NDRG1), which is involved in suppression of metastasis, particularly under hypoxic conditions (Salnikow et al., 2002; Mao et al., 2013).

#### The spleen

Uniquely abundant transcripts in the spleen tended to encompass more functional diversity than transcripts identified for the other tissues sampled. Highly abundant spleen-specific transcripts include proteins involved in nucleotide exchange (ARHGEF17, TPM= 137.25), erythropoiesis (EPOR TPM= 26.56, SPTA1 TPM= 21.82), and GTP hydrolysis (GBP6, TPM= 22.15), as well as at least one kinase (LIMK2, TPM= 10.09) that is associated with immune function (Bernard, 2007; Kim et al., 2011; Lutz et al., 2013; Ponceau et al., 2015; Kuhrt & Wojchowski, 2015). Both erythrocytic activity and immune function are consistent with the functional role of the spleen, which filters blood and recycles blood cells (Cesta, 2006; Scott & Olson, 2007; Droppelmann et al., 2013; Pivkin et al., 2016). Interestingly, the spleen was found to express more genes under positive selection than expected (Fig. 1), suggesting this tissue may be an active target for adaptation. Three of these genes (Sperm Associated Antigen 9; SPAG9, Cell Division Cycle 7; CDC7, and Zinc Finger CCCH-Type Containing 13; ZC3H13) have been previously characterized in humans. Upregulated in cancerous cells, SPAG9 is thought to be an early marker for diagnosis (Baser et al., 2013; Chen et al., 2014). Cell Division Cycle 7 is a DNA replication regulator, and can inactivate tumor suppressor protein p53 when CDC7 is overexpressed in tumor cells (Bonte et al., 2008; Ito et al., 2012). Finally, ZC3H13 is a component of Wilms’ tumor associating protein, a splicing regulator potentially required for cell cycle progression (Horiuchi et al., 2013).

#### The liver

The primary functions of the liver are to produce blood coagulation hormones, detoxify blood, and to metabolize foreign substances (Cheeke, 1994; Wada, Usui & Sakuragawa, 2008; Davidson, Ballinger & Khetani, 2016; Schiöth et al., 2016; Harrall et al., 2016). The two genes that were most uniquely expressed in the liver were associated with these functions, specifically blood clotting (Fibrinogen Alpha Chain: FGA, TPM= 864.02), and drug toxin metabolism (Cytochrome P450 2A11: CYP2A11, TPM= 370.43) (Mosesson, 2005; Yang et al., 2012). Our results suggest that the liver, like the spleen, may also be an active site of adaptation given the number of genes found to be under positive selection in the liver was more than twice that expected by chance (Fig. 1). Of these genes, three are involved in metal ion transport (Solute Carrier Family 30 Member 10 [SLC30A10], Nedd4 Family Interacting Protein 2 [NDFIP2], and Family With Sequence Similarity 21 Member C [FAM21]) (Ohana et al., 2006; Yang et al., 2012; Shusterman et al., 2014; Gallon & Cullen, 2015; Lee, Chang & Blackstone, 2016; Foot et al., 2016), while three others have ontology terms associated with immune response (Signal Peptide Peptidase Like 2A [SPPL2A: Biological Process-regulation of immune response], Ataxin 2 [ATXN2: Biological Process - negative regulation of multicellular organism growth], SET Domain Containing 6 [SETD6: Biological Process-regulation of inflammatory response]). Given the roles that the spleen and liver play in immunological processes and the genes identified to be under positive selection in these tissues, it is possible that both the spleen and liver of the tuco-tuco are particularly involved in adaptation to the subterranean environment.

*C. sociabilis* is not the first subterranean rodent to provide evidence of possible adaptation to the regulation of cell cycling. The naked mole-rat (*H. glaber)*, has been the subject of numerous studies attempting to discern the source of the cancer resistance reported for this long-lived species (Buffenstein, 2008; Rodriguez et al., 2011; Delaney et al., 2013). Decreased prevalence of cancer in the naked mole rat has been attributed to a heightened sensitivity to contact inhibition (Seluanov et al., 2009) and fibroblast secretion of high-molecular-mass hyaluronan (Tian et al., 2013). Studies have also suggested that the naked mole rat has increased translational fidelity due to a unique 28S ribosomal structure (Azpurua et al., 2013). More recently, cancer has been detected in this species (Delaney et al., 2016), although these examples were based on studies of captive mole-rats not exposed to the natural hypoxic environment for this species, an environmental setting that may have contributed to tumor formation (Welsh & Traum, 2016). Colonial tuco-tucos also presumably occur in hypoxic environments and it is possible that the fourteen apoptotic genes identified as being subject to positive selection in this species also have important regulatory functions in this setting. Gene ontology terms associated with cell cycling/DNA damage response genes comprised over 20% (12 genes of 50) of the genes identified as being under positive selection, with other gene ontology categories comprising a substantially smaller portion of the loci thought to be subject to selection. Collectively, these genes present important candidates for future studies of regulation of cell physiology in subterranean rodents.

Future studies of *C. sociabilis* would benefit from quantifying differential gene expression across multiple individuals to provide a more robust quantitative assessment of tissue-specific patterns of gene expression. Of the highly abundant transcripts identified for each tissue type, many suggest a role in immune function while positively selected genes hint at specializations for cell cycle regulation. Both of these characteristics are seen across the different tissues samples for *C. sociabilis*. Expression patterns can vary greatly among individuals, and thus although our data set does not allow for statistical analyses of patterns of gene expression in *C. sociabilis*, our findings are consistent with those revealed by previous studies of subterranean organisms. Expansion of our analyses to include multiple individuals, as well as additional taxa, will allow for a more comprehensive understanding of the genomic underpinnings of physiological adaptations to subterranean life.

## SUMMARY

In this study, we present a high quality and complete transcriptome for the colonial tuco-tuco (*C. sociabilis)*. By characterizing transcriptomes generated from eight tissue types, we provide preliminary insights into how transcript abundance differs across tissues. Notably, the most abundant transcripts and the genes subject to positive selection were generally consistent with the primary physiological function(s) of the tissues from which they were derived, with a prevalence of transcripts associated with cell proliferation. We also identify a set of genes that appear to be under positive selection; the number of genes subject to selection that were expressed in the liver and spleen were greater than expected, suggesting that these tissues are of particular functional importance to the colonial tuco-tucos. The underlying reasons for enhanced selection of genes in these tissues remains to be determined, providing an intriguing basis for additional studies of genomic evolution in *C. sociabilis* and other subterranean rodents. At the same time, given extensive field data regarding the behavior, ecology, and physiology of *C. sociabilis*, the transcriptomic data presented here represent a critical tool for future studies aimed at clarifying relationships among physiology, selection, and specialization for a subterranean lifestyle.

